# Pre-stimulus alpha activities predict the confidence in subjective judgment by modulating post-stimulus theta oscillation

**DOI:** 10.1101/2021.01.19.426904

**Authors:** Haoyue Qian, Yang Lu, Zhiyuan Liu, Xue Weng, Xiuyan Guo, Lin Li, Li Zheng, Min Xu, Xiaohan Cai, Weiwei Men, Jiahong Gao, Xiangping Gao

## Abstract

Even when making arbitrary decisions, people tend to feel varying levels of confidence, which is associated with the pre-stimulus neural oscillation of the brain. We investigated varying confidence in a pure subjective judgment task, and how this confidence was predicted by pre-stimulus alpha oscillations. Participants made pure subjective judgments where their prior experience seems to be helpful but actually useless, and their fluctuating confidence was related to the choice boundary process rather than the evidence accumulation process, suggesting participants underwent varying confidence resulting from the internal signals. With EEG and MEG analyses, we not only revealed the linkage between confidence and pre-stimulus alpha activities, but also successfully located this linkage onto decision-making relevant brain areas, i.e. MCC/PCC and SMA. Moreover, we unveiled a specific pathway underlying such linkage, that is, the influence of pre-stimulus alpha activities on decision confidence was fulfilled through modulating post-stimulus theta activities of SMA.

## Introduction

People make decisions and undergo the sense of confidence, not only when decisions are based on the process of external information, but also when there are ambiguous or insufficient sensory evidence and therefore the decisions are substantially determined by internal states. Previous studies investigated such decisions in tactile and auditory perceptual discrimination tasks when the physical signals from stimuli are constant, and revealed that, decision confidence was fluctuated and predicted by pre-stimulus brain internal state especially indicated by neural oscillations in alpha band (Baumgarten et al., 2016; Woestmann et al., 2019). Still, it remains unclear which pathway supports the modulation of pre-stimulus alpha oscillation on decision confidence experienced after stimulus, which is the main concern of the current study.

Numerous studies have showed that low frequency neural activities, most prominently in the alpha-band (8 ~ 12Hz), occur spontaneously within many brain areas and have been linked to the following cortical excitability (Haegens et al., 2011; Harvey et al., 2013; van Kerkoerle et al., 2014). Recent research clearly revealed that during perceptual decision, the pre-stimulus alpha power was relevant to the post-stimulus neuronal excitability associated with participants’ decision responses (Romei et al., 2010; Lange et al., 2013; Chaumon & Busch, 2014). Taking these evidences together, we inferred that pre-stimulus alpha oscillations influence the subjective decision through modulating the post-stimulus neuronal activities.

A recent study addressed the issue but failed to find the influence of pre-stimulus alpha power on decision confidence is related to the post-stimulus neural activities (Woestmann et al., 2019). A possible account for the failure of this study to identify the potential influence pathway is that the researchers only focused on the neural activities of sensory brain areas in perceptual decision. Note that the influence of spontaneous neural oscillations in alpha band on the following cortical excitability occur in a variety of brain areas except for sensory areas (Haegens et al., 2011; van Kerkoerle et al., 2014). In addition, data-driven analyses in recent EEG studies showed the effect of pre-stimulus neural oscillation on subjective decision was widely distributed over many electrodes (Iemi et al., 2017; Benwell et al., 2017). Given the fact that the pre-stimulus neural oscillation could predict the varying confidence even in the absence of external sensory evidence (Woestmann et al., 2019), the critical brain areas associated with the influence of pre-stimulus alpha power on confidence may be other than sensory areas. For instance, they could be such as SMA (supplementary motor area) that directly contributes to the processing of decision confidence (Fleming et al., 2015), and PCC (posterior cingulate cortex) that has been implicated in subjective valuation of the decision error (Bach et al., 2011). To investigate whether some decision-making relevant brain areas may support such influence, we explored the relevant areas at whole-brain level with MEG and EEG adopting a subjective judgement task.

Here we recruited Chinese participants to make pure subjective judgments, i.e., to assess the Jurchen word-likeness for unfamiliar stimuli, of which, though, some sub components may be experienced before. That is to say, their prior experience seems to be helpful but was actually useless in the task. Besides EEG, MEG recordings were also used to observe pre-stimulus neural oscillation and post-stimulus neural activities with high temporal and spatial resolution. With the help of PTE (phase transfer entropy) method, it is feasible to achieve a clear explanation on how pre-stimulus neural activities of specific brain areas (e.g., SMA and PCC) and their interplay could modulate the post-stimulus neural activities, and finally influence confidence.

## Results

### The confidence was fluctuating during subjective judgment

In EEG and MEG experiments, participants, who are native Chinese speakers and have no experience on Jurchen, were separately recruited to judge whether the stimuli were real or pseudo Jurchen character with different confidence choices (low vs. high confidence; Fig. 1a, details in method). We found all participants focused on the task according to their relatively high rates of responding correctly to catching trials that instructed participants to press a specific button (for each participant, the rate was higher than 83.33%).

**Figure 1.**
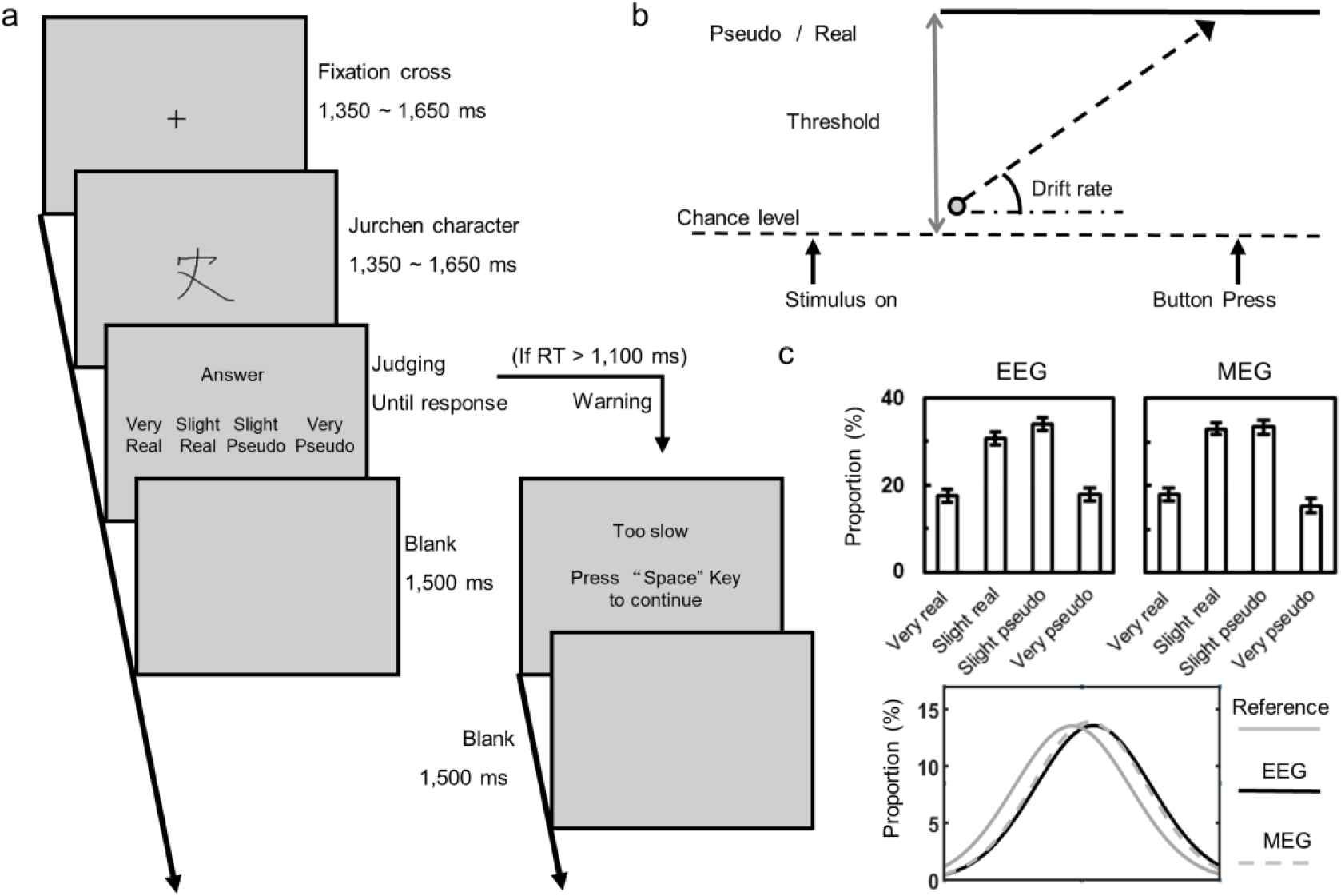
Experimental design and stimuli. a. An example trial for the subjective judgement task. Participants saw the Jurchen character stimulus for 1,500 ms, following a 1,350-1,650 ms fixation cross. Then, they judged whether the stimulus was a real or pseudo Jurchen character with four choices, i.e. very real, slightly real, slightly pseudo, and very pseudo. If participants failed to give a response within 1,100 ms, the trial terminated with a warning message and they pressed ‘Space’ key to continue. A new trial began after a 1,500 ms blank screen. Of note, all stimuli were real Jurchen characters and all texts were present in Chinese. b. The drift diffusion model of the subjective judgment task. It is assumed that each decision is an accumulation of noisy information indicative of a true or pseudo choice. After the onset of stimulus, relative evidence starts to accumulate from (around) the chance level up to the boundary (i.e. one of the choices) and finally initiates the corresponding response. The model contains two key parameters, i.e. drift rate and threshold. The drift rate is defined as the speed of the accumulation, representing processes of evidence accumulation. The threshold is defined as the distance between the two boundaries, representing processes of decision boundary. c. Frequency distributions of choices and distribution fitting. The frequency distributions of four choices based on EEG and MEG data consistently showed, there were about twice as many low confidence choices as high confidence choices (upper). We fitted normal curves to the frequency distributions of choices with different confidence levels based on our EEG and MEG experimental data as well as the data from the study of Woestmann and colleagues (i.e. Reference; Woestmann et al., 2019) respectively (bottom). This resulted in three fitting curves that were very similar. Error bars represent s.e.m.

To examine how participants underwent the sense of confidence in this subjective task, we tested whether there was a pattern in the distribution of participants’ confidence choices. As Figure 2a illustrated, in EEG experiment, the ratio of low confidence choice (M = 64.64%) was higher than that of high confidence choice (M = 35. 36%), *V* = 18, *P* < 0.001, 95% CI = 20.00% to 40.42%; in MEG experiment, the ratio of low confidence choice (Low confidence, M = 66. 45%) was also higher than that of high confidence choice (High confidence, M = 33.55%), *V* = 14.5, *P* < 0.001, 95% CI = 23.33% to 41.67%. The RT analyses showed, the RT of low confidence trials was longer than that of high confidence in both EEG experiment, *t*(27) = 5.23, *P* < 0.001, Cohen’s d = 0.99, 95% CI = 36.80 to 84.29, and MEG experiment, *t*(29) = 8.09, *P* < 0.001, Cohen’s d = 1.48, 95% CI = 46.76 to 78.38 (Fig. 2b). The results suggested, participants actually underwent varying levels of confidence, and the varying confidence accompanied with variance of RT, lower confidence to longer RT.

**Figure 2.**
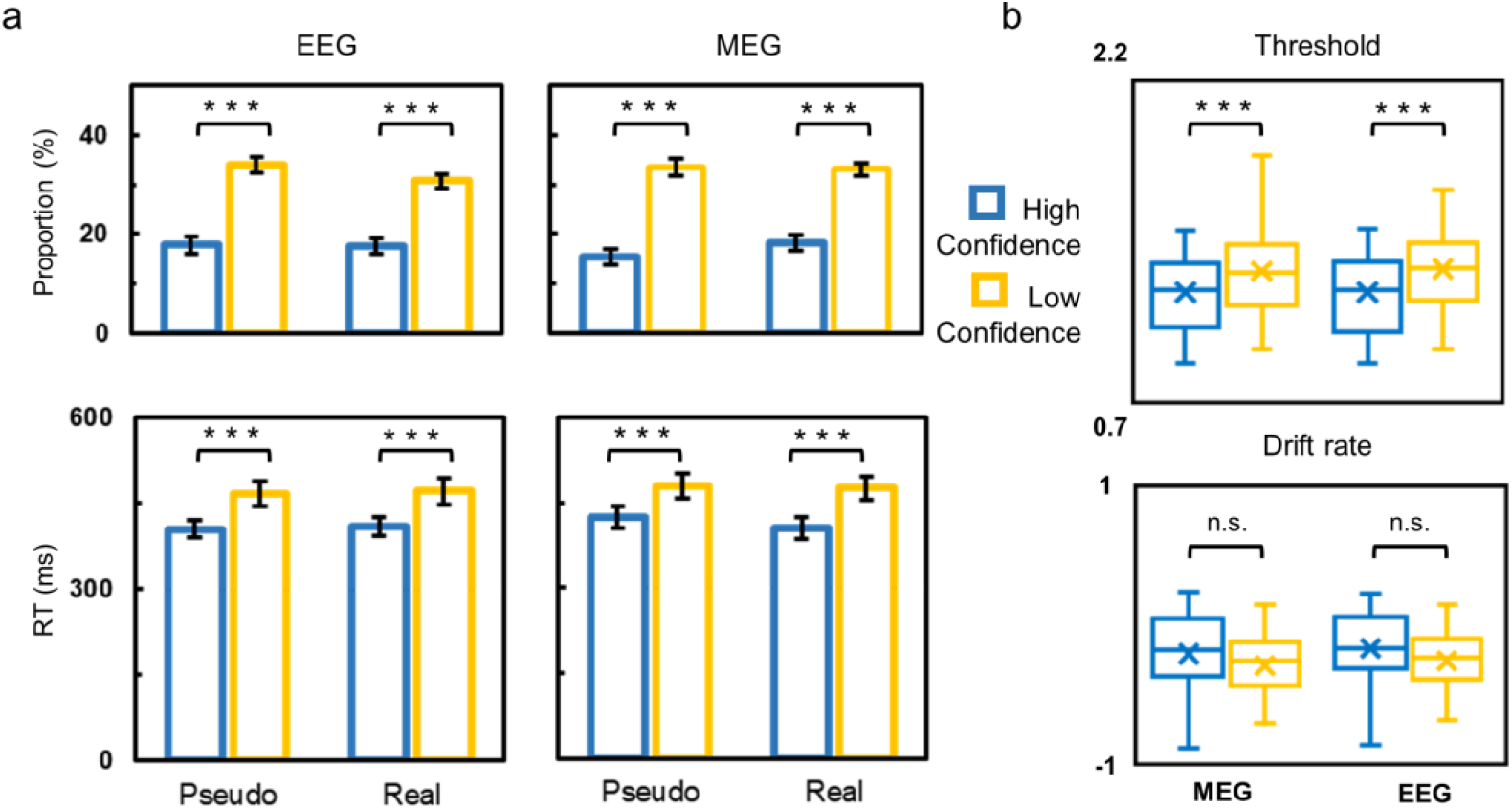
The behavioral results and model parameters in the subjective judgment task. a. The proportions and RTs of four choices in EEG (left) and MEG experiments (right). In both experiments, participants had lower proportion of choices and shorter RTs in the high confidence choice trials than in low confidence choice trials. b. The drift diffusion model fitting parameters for the data of each participant. In both experiments, the significant difference between low and high confidence trials was found in thresholds but not in drift rates. The orange represents the data from low confidence choice trials while the blue represents the data from high confidence choice trials. The box represents the interquartile range (IQR). The line through the middle is the median. Whiskers extend to 1.5 × IQR; outliers are plotted as dots. \*\*\**P* < 0.001. Error bars represent s.e.m.

Interestingly, the ratio of high confidence choice in this study (EEG: 35.36 %; MEG: 35.55%) was very similar with that of a previous study which focused on perceptual decision driven purely by internal signals (Woestmann et al., 2019: 38.10%; Fig. 1c). This fact was consistent with the notion that participants' confidence in this subjective judgement task might be mostly associated with their internal signals.

Additionally, according to drift-diffusion model (DDM), we examined whether the varying confidence in this subjective judgment was shaped by the processes of evidence accumulation, or boundary, or both. To address this, we fitted DDM to behavioral data for each participant, and estimated parameters of drift rate and threshold in high and low confidence trials respectively (Fig. 1b). According to DDM model, drift rate and threshold reflect the process of evidence accumulation and boundary respectively. The paired t test on the thresholds showed significant difference between high and low confidence trials in both EEG experiment, *t*(27) = 4.52, *P* < 0.001, Cohen’s d = 0.85, 95% CI = 0.06 to 0.16, and MEG experiment, *t*(29) = 5.20, *P* < 0.001, Cohen’s d = 0.95, 95% CI = 0.08 to 0.17. However, we did not find the significant difference in drift rates between high and low confidence trials in either EEG experiment, *t*(27) = 0.48, *P* = 0.632, 95% CI = −0.27 to 0.16, or MEG experiment, *t*(29) = 1.93, *P* = 0.064, 95% CI = −0.52 to 0.02. Thus, the confidence of this task was related to the choice boundary process and relevant internal signals, rather than the evidence accumulation process driven by external inputs.

In short, the behavioral results suggested, when making this subjective judgement, participants still underwent the sense of confidence and this confidence depended on the internal signals.

### The pre-stimulus alpha oscillations of SMA and MCC/PCC were associated with the confidence in subjective judgment

Having demonstrated that participants underwent varying confidence, which was associated with the internal signals, we next investigated whether spontaneous (i.e. pre-stimulus) brain activities were correlated with the confidence.

To address this, we first plotted the time frequency map of contrast (low vs. high confidence) (Fig. S1). The maps of EEG and MEG consistently suggested the pre-stimulus neural oscillation related to the confidence was in low alpha band (around 8 ~ 9 Hz) from 1000 to 500 ms before stimulus onset. Then, we explored the topographical location and cortical source of the confidence-related pre-stimulus oscillation. The EEG topography suggested that the alpha oscillation was mainly from the central scalp region, C2 and C4, and the MEG topography indicated two sensor-clusters at central scalp region for the alpha oscillation (Fig. 3a). The source reconstruction with MEG data identified two cortical sources for the pre-stimulus alpha oscillation (*P* < 0.02, cluster corrected, Fig. 3b), i.e. MCC/PCC (medial and posterior cingulate cortex; MNI: 1.2, −14, 37) and SMA (supplementary motor area; MNI: −3.4, −15, 57). Finally, PSD (power spectral density) map clearly demonstrated in EEG, pre-stimulus low alpha power at central scalp region was higher in low confidence trials comparing to high confidence trials (*P* < 0.05); in MEG, the pre-stimulus low alpha power of MCC/PCC and SMA was higher in low confidence trials comparing to high confidence trials (*P* < 0.01). Together, by comparing the low and high confidence trials, we found the pre-stimulus alpha power was higher in low confidence trials (vs. the high confidence trials).

**Figure 3.**
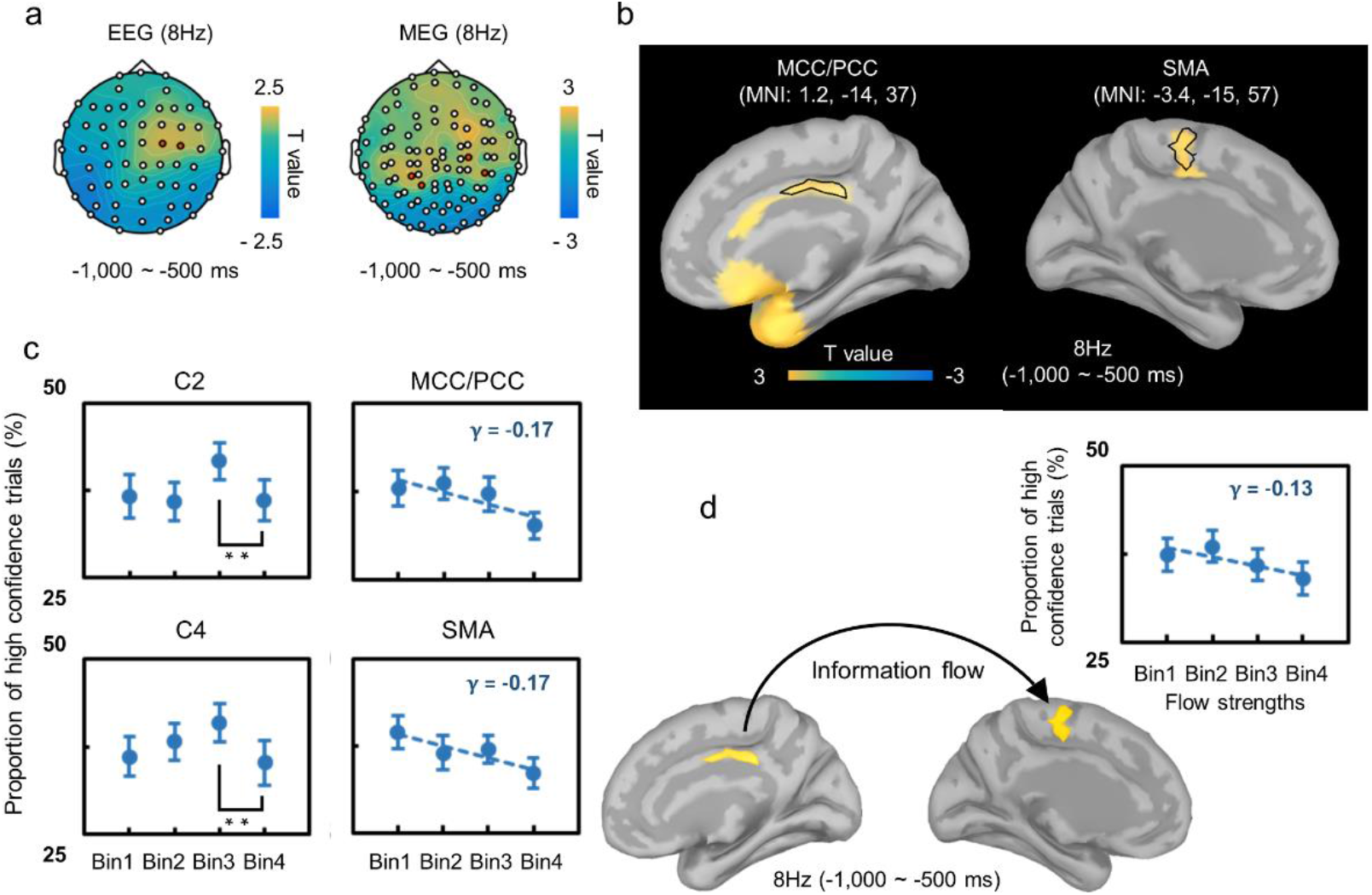
The pre-stimulus neural activities associated with the confidence in subjective judgment. a. the t-value topographies of low vs. high confidence trials. In EEG experiment, the alpha (8 Hz) oscillations were from the central scalp region, electrodes C2 and C4 (red-marked electrodes); In MEG experiment, the alpha oscillations were observed in two sensor clusters in central scalp region (red-marked sensors). b. MEG source localization results of the pre-stimulus alpha oscillation identified the two brain areas, MCC/PCC and SMA (*P* < 0.02, cluster corrected). c. Group-averaged changes in high confidence choice ratios according to pre-stimulus alpha powers of EEG (left) and of MEG (right). The data from C2 and C4 showed the high confidence choice ratio in the bin 4 (i.e. the strongest) was significantly lower than that in the bin 3, suggested the potential relation between the high confidence choice ratios and the pre-stimulus oscillations. The data of MCC/PCC and SMA further confirm the relation, showing that the high confidence choice ratio decreased along the pre-stimulus alpha power bin ranks of MCC/PCC and SMA. d. Information flow strength of the pre-stimulus alpha oscillation from MCC/PCC to SMA, as indicated by the phase transfer entropy analyses, was correlated to the ratios of high confidence choices. \*\**P* < 0.01. Error bars represent s.e.m.

We further investigated whether the pre-stimulus low alpha power predicted the probability of high confidence choice. For each trial per participant, we computed the low alpha power (i.e. 8Hz) averaged over the time window from −1000 to −500 ms. For each participant, trials were sorted from low to high power and divided into 4 bins (the number of trials was equal in each bin), and then, each participant's ratio of high confidence choice in each of 4 bins was obtained respectively by computing the proportion of trials with high confidence choice in each bin. We adopted a random-intercept-fixed-slope multilevel model to explore whether the low alpha power of pre-stimulus neural oscillation predicted the ratio of high confidence choice.

The model did not show the ratios of high confidence choice varied along the bin ranking of pre-stimulus alpha power at C2, *γ* = 0.032, *t*(23) = 0.54, *P* = 0.596, 95% CI = −0.09 to 0.15 or C4, *γ* = −0.001, *t*(23) = 0.01, *P* = 0.997, 95% CI = −0.11 to 0.11. Nevertheless, the repeated measures ANOVA showed, the ratios of high confidence choice in four bins were significantly different [C2: *F*(3,69) = 3.37, *P* = 0.023, partial*η*^2^ = 0.13; C4: *F*(3,69) = 3.71, *P* = 0.015, partial*η*^2^ = 0.14]; the high confidence choice ratio in the fourth power bin (i.e. the strongest one) was significantly lower than that in the third power bin [C2: *t*(23) = 3.14, *P* = 0.027 (Bonferroni corrected), Cohen’s d = 0.64, 95% CI = 0.019 to 0.092; C4: *t*(23) =3.10, *P* = 0.030 (Bonferroni corrected), Cohen’s d = 0.63, 95% CI = 0.019 to 0.096, suggesting a possibility that there was a negative co-variation between the pre-stimulus alpha power and the high confidence ratio. The negative co-variation was clearly confirmed by MEG data. The model revealed the high confidence choice ratio decreased along the pre-stimulus alpha power bin ranks of SMA, *γ* = −0.17, *t*(72) = 3.16, *P* = 0.002, Cohen’s d = 0.37, 95% CI = −0.27 to −0.06, as well as those of MCC/PCC, *γ* = −0.17, *t*(53) = 2.90, *P* = 0.006, Cohen’s d = 0.40, 95% CI = −0.29 to − 0.05 (Fig. 3c). The MEG results demonstrated that the pre-stimulus low alpha power of MCC/PCC and SMA could predict the probability of high confidence choice in subjective judgment.

In addition, we investigated the directional information flow between these two brain areas in alpha band based on MEG data. To address this, we used phase transfer entropy (PTE, see method) to calculate the information flow strength between MCC/PCC and SMA at 8 Hz within pre-stimulus period (−1000 ms to −500 ms). We applied the normalized strength of information flow, ranging from −0.5 to 0.5. The positive value indicated the information flow from MCC/PCC to SMA and the negative value indicated the information flow from SMA to MCC/PCC. The analysis revealed, the strength of information flow in low confidence trials was significantly greater than zero (0.0103, one tailed *t*(25) = 1.83, *P* = 0.034, Cohen’s d = 0.36), whereas the information flow strength in high confidence trials was not significantly different from zero (−0.0046, one tailed *t*(25) = 0.80, *P* = 0.216). The permutation one-tailed Wilcox test further confirmed that the information flow strength was significantly larger in low confidence trials comparing with that in high confidence trials, *W*(26) = 145, *P* = 0.035. Besides, binning analysis showed, high confidence choice ratio decreased along the bin ranks (from low to high) of the information flow strength from MCC/PCC to SMA,*γ* = −0.126, *t*(25) = 2.23, *P* = 0.035, Cohen’s d = 0.44, 95% CI = −0.24 to −0.01 (Fig. 3d). These results revealed the pre-stimulus MCC/PCC-to-SMA information flow of alpha band in low confidence trials but not in high confidence trials, and the strength of information flow was negatively related to the probability of high confidence choice.

In short, the pre-stimulus MCC/PCC-to-SMA information flow in alpha band could predict the variance of confidence in subjective judgment.

### The prediction of pre-stimulus alpha oscillations on confidence was mediated by post-stimulus theta oscillation of SMA

After revealing the influence of pre-stimulus neural activities of MCC/PCC and SMA on the confidence, we sought to investigate the potential pathway underlying the influence. According to previous studies (Palva & Palva, 2007; Tenke et al., 2015; Iemi et al., 2019), it is probable that the influence of pre-stimulus alpha power on following neural activity occurs in the same brain area. We first identified the post-stimulus neural oscillation of SMA and MCC/PCC for low vs. high confidence contrast. Then, we examined whether the prediction of pre-stimulus activity on the confidence was mediated by the post-stimulus neural oscillation.

As Figure 4a showed, the EEG time-frequency map of the central scalp region (C2 and C4; *P* < 0.02) indicated the powers of post-stimulus were significantly larger for low confidence choices compared with high confidence choices oscillation at three frequencies. The first one was around 4 Hz from 1650 to 2200 ms after stimulus onset; the second one was around 9 Hz from 650 to 1050 ms after stimulus onset; the third one was around 14Hz from 1350 to 1550 ms. The low vs. high confidence contrast for post-stimulus oscillation in MEG data was very similar with that in EEG.

**Figure 4.**
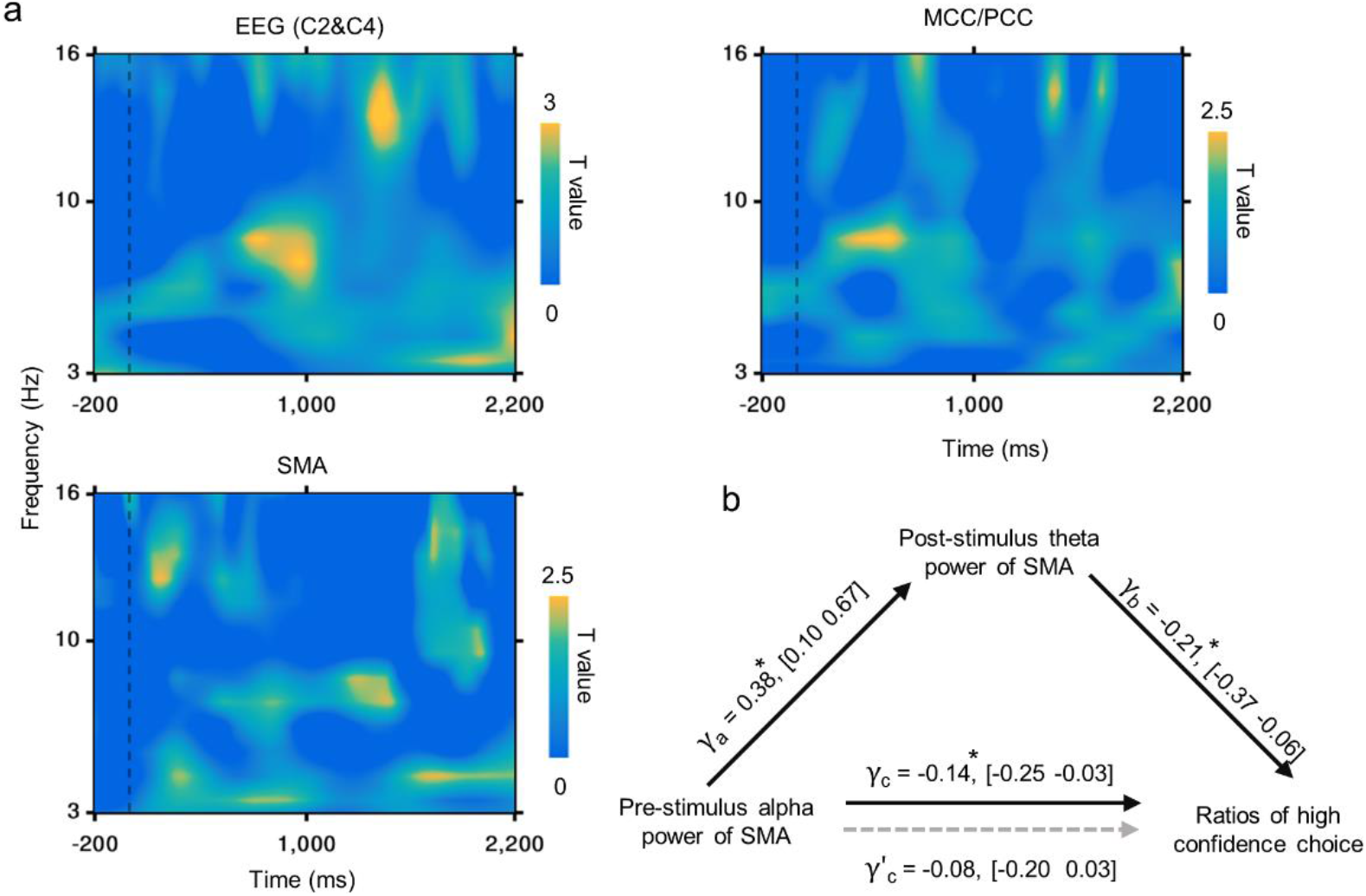
The post-stimulus neural oscillations related to the confidence and its mediating effect. a. Time-frequency t-value map of low vs. high confidence trials. The t-value map of EEG at the central scalp region (C2 and C4; *P* < 0.05) indicated the powers of post-stimulus oscillation were significantly larger for low confidence choices compared with high confidence choices at three frequencies. The first one was around 4 Hz from 1,650 to 2,200 ms after stimulus onset; the second one was around 9 Hz from 650 to 1,050 ms after stimulus onset and the third one was around 14Hz from 1,350 to 1,550 ms (upper left). The low vs. high confidence contrast for post-stimulus oscillation in MEG data was very similar with that in EEG. The map of SMA (*P* < 0.05) revealed confidence-related theta oscillation (5 Hz) from (bottom left) and the map of MCC/PCC revealed significant alpha oscillations (9 Hz) from 250 to 600 ms (upper right). b. The prediction of pre-stimulus alpha oscillations in SMA on confidence was mediated by its post-stimulus theta oscillation. After controlling the mediation effect of the post-stimulus theta oscillations, the regression of pre-stimulus alpha oscillations to the ratios of high confidence choice became not significant. \**P* < 0.05. The values in square brackets are 95% confidence intervals.

Specifically, the time-frequency map of SMA (*P* < 0.05) revealed the theta oscillation (5 Hz) from 1600 to 1900 ms was significantly larger for low confidence choices than high confidence choices, and that of MCC/PCC revealed the alpha oscillation (9 Hz) from 250 to 600 ms was significantly larger for low confidence choices than high confidence choices.

Before we conducted the mediation analysis for the effect of pre-stimulus neural activities on high confidence ratios, we need to confirm the significant relation between pre-stimulus neural activities and post-stimulus neural oscillation. To test the relation, we applied binning analysis based on MEG data (see method). The analysis revealed a positive correlation between the pre-stimulus alpha power and the post-stimulus theta power in SMA, *γ* = 0.38, *t*(26) = 2.78, *P* = 0.010, Cohen’s d = 0.54, 95% CI = 0.10 to 0.67, whereas the correlation between the pre-stimulus alpha power and the post-stimulus alpha power in MCC/PCC was not significant,*γ* = 0.22, *t*(26) = 1.49, *P* = 0.148, 95% CI = −0.08 to 0.53. Besides, the analysis showed, the correlation between the information flow strength from MCC/PCC to SMA and the post-stimulus theta power of SMA was not significant, *γ* = 0.04, *t*(25) = 0.34, *P* = 0.736, 95% CI = −0.18 to 0.25.

To identify the potential mediation role of post-stimulus theta oscillation of SMA in the effect of its pre-stimulus alpha oscillation on the confidence, we examined whether the correlation between pre-stimulus alpha power and confidence would substantially decrease under statistical control for the influence of post-stimulus neural oscillation. The binning analysis suggested, after excluding variance in high confidence ratios explained by the post-stimulus theta power of SMA, the correlation between pre-stimulus alpha power and high confidence ratio became not significant (from *γ* = −0.14, *P* = 0.011 to*γ* = −0.08, *P* = 0.165; Fig. 4b). The results above revealed the influence pathway of pre-stimulus alpha oscillation in SMA on high confidence ratios, i.e. such influence was entirely mediated by the post-stimulus theta oscillation in SMA.

In short, when participants made subjective judgement, their varying confidence was predicted by pre-stimulus alpha activities of SMA and PCC, and this prediction was mediated by post-stimulus theta oscillation in SMA.

## Discussion

In this study, we applied a pure subjective judgement task with EEG and MEG recordings to examine how pre-stimulus neural oscillation influenced post-stimulus neural activities and finally the decision confidence. First, analyses on behavioral data revealed, in subjective judgement task, participants reported fluctuating confidence, and the distribution of confidence kept consistent with that of perceptual decision driven purely by internal signals (Woestmann et al., 2019). Converging findings from DDM analyses proved the confidence was related to the choice boundary process rather than the evidence accumulation process. These suggested participants underwent the sense of confidence during subjective judgement task and their confidence depended on the internal signals. Then, we explored how specific brain areas and their interplay involved in the influence of pre-stimulus alpha oscillations on confidence. Pre-stimulus MCC/PCC-to-SMA information flow were found as associated with confidence, which means varying levels of confidence was predicted by not only the pre-stimulus alpha oscillations of MCC/PCC and SMA, but also the information flow strength in alpha band from MCC/PCC to SMA. Finally, with a mediation analysis we addressed the linkage between pre-stimulus alpha oscillations and confidence, i.e., to identify which specific post-stimulus neural activities were modulated during the influence of pre-stimulus alpha oscillations on confidence. We found that pre-stimulus alpha activities of SMA, which receiving information flow from MCC/PCC, influenced confidence through variation in its own post-stimulus theta oscillations.

Contrast to previous study that focused on the pre-stimulus neural activities in sensory areas (Woestmann et al., 2019), we thoroughly explored the brain areas related to the influence of pre-stimulus alpha oscillations on confidence at whole-brain level with both MEG and EEG. Such a data-driven approach comparing the low and high confidence trials identified MCC/PCC and SMA as relevant brain areas. Using binning analyses, we confirmed that the pre-stimulus powers of MCC/PCC and SMA could predict the variance of participants’ confidence. Research showed, in decision task, PCC engages in subjective estimation of the probability of error (Bach et al., 2011) and uncertainty (Paul et al., 2015). Considering that participants need to make subjective judgements with uncertainty in the current study, the pre-stimulus activities of MCC/PCC found here can be considered as the spontaneous activities that related to the following subjective estimation of uncertainty and thereby confidence. As to SMA, it is a region playing an important role in response or decision threshold setting that affected trial-by-trial decision (Forstmann et al., 2008; Forstmann et al., 2010). Some follow-up studies using DDM also revealed SMA functioned as tracking the variation of choice boundary (Georgiev et al., 2016), and further influenced the varying levels of confidence (Bang & Fleming, 2018). Therefore, in the current study, the activities of SMA may directly contribute to subjective decision threshold setting and confidence estimating (Fleming et al., 2015). Besides, in the current study, the PTE analyses revealed the pre-stimulus MCC/PCC-to-SMA information flow in alpha band. Previous research found SMA has a functional connectivity with PCC at resting state, and such connectivity was associated with specific behavioral patterns (Zhang et al., 2016). Consistent with the finding, here the current study suggested the connectivity between SMA and MCC/PCC, indexed by strength of information flow, might indicate a confidence-related behavioral pattern. Note that, as an important node of default mode network (DMN; Hagmann et al., 2008; van Oort et al., 2014), PCC has found to be a strong driver of spontaneous activity in healthy humans (Coito et al., 2019). Our PTE result was then in line with these findings, suggesting MCC/PCC worked as an originator of the decision-confidence related internal signals in subjective judgement and transferred the information in alpha band to SMA.

In order to identify the specific pathway that the pre-stimulus oscillation affects confidence in post-stimulus decision-making, we examined whether the pre-stimulus alpha activities influenced confidence by modulating the post-stimulus neural activities. In the post-stimulus time window, we first found theta oscillations of SMA and alpha oscillations of MCC/PCC were associated with confidence. Nevertheless, binning analyses revealed the only significant correlation between the pre-stimulus alpha oscillations of SMA and its own post-stimulus theta oscillations, suggesting SMA as the sole brain region that the potential modulation of pre-stimulus neural activities on the post-stimulus neural activities occurred. Follow-up mediation analyses confirmed that the influence of pre-stimulus neural oscillations of SMA on confidence was fully mediated by its own post-stimulus neural activities. Combining the finding that SMA involved in choice boundary (Bang & Fleming, 2018) with our finding that the confidence in this subjective judgement task was shaped by the process of choice boundary, the mediation findings above indicated that the pre-stimulus alpha activities modulated post-stimulus neural activities associated with boundary process, and thereby the confidence. As documents illustrated, the spontaneous neural activities in low bands could modulate the cortical excitability thereafter (Haegens et al., 2011; Harvey et al., 2013; Jensen & Mazaheri, 2010). In perceptual decision, it has been established that pre-stimulus neural activities affect the post-stimulus neural activities (Romei et al., 2010; Lange et al., 2013; Chaumon & Busch, 2014). Extending this line of studies, we found in the present subjective judgement task, it was SMA where the pre-stimulus neural activities influenced decision confidence by modulating the post-stimulus neural activities. Hence, our findings supported such a general mechanism that the spontaneous neural oscillations in low bands influenced following neural activities could be widely adopted to explain connection between the pre-stimulus neural activities and subsequent decision confidence.

Taken together, the current study demonstrated a complete influence pathway of pre-stimulus neural oscillations on confidence in subjective judgment, i.e. MCC/PCC served as an originator of the internal signals before stimulus onset and transferred the information in alpha band to SMA; after that, the pre-stimulus alpha activities of SMA modulated its post-stimulus theta activities and relevant boundary process, and finally the confidence of subjective judgment.

## Method

### Participants

Twenty-eight healthy adults (16 females; mean age, 23.57 ± 2.03y) were recruited for the EEG experiment and another thirty adults (14 females; mean age, 21.03 ±1.85y) were recruited for the MEG experiment. The sample size ensured us to indicate the significant (α = 0.05, two tails) influence of the pre-stimulus alpha power on subjective confidence (Woestmann et al., 2019; effect size: Cohen’s d = 1.28) or on post-stimulus neural activities (Iemi et al., 2019; effect size: Cohen’s d = 1.05), given that the power is set at 0.95. All participants were right-handed, native speakers of Chinese with no history of neurological nor psychiatric disorders. None of participants had experience for the Jurchen script. Prior to experiment, all participants gave informed consent, and after participating, they received the compensation (RMB 60/h). The EEG and MEG experiments were approved by the Ethics Committee from the universities where the data were collected, respectively.

### Materials

In the task, all character stimuli were real characters in the Jurchen dictionary (Jin, 1984). Jurchen is an abandoned non-alphabetic writing language (Kane, 1975). The structures and sub components of the Jurchen characters were very similar to the Chinese characters (Shen, 2011). To select the character stimuli without obvious external features for Chinese participants, we excluded the Jurchen characters with too many (over 10) or too few strokes (below 4) and those having same sub components as the Chinese characters. Besides, based on the Chinese character likeness ratings with a 5-point Likert scale (from very high likeness to very low likeness) from 10 adults who did not participate the formal test, we also excluded the Jurchen characters that were too similar to Chinese characters (below 2.5 out of 5) as well as those were extremely dissimilar to Chinese characters (above 4.5 out of 5). Finally, 280 were selected from 859 Jurchen characters, and these selected characters had low likeness level Chinese characters (M =3.78, SD = 0.45). Thus, the participants’ Chinese character experience seems to be helpful but actually is useless. Among the 280 characters, 40 were randomly selected as a pool for demo and practice and the remaining 240 were used for test.

### Procedure

On each trial, participants were asked to focus on a fixation cross (1° visual angle) during a random length (1350 ms – 1650 ms). Then, the Jurchen character stimulus (3.9° visual angle) was presented for 1500 ms, followed by the response screen. Referring to previous research did on subjective judgement with uncertainty (Colas & Hsieh, 2014; Horr et al., 2014), we required participants to give responses within 1100ms, judging whether the stimulus presented was a real Jurchen character or not among the four choices provided. When participants failed to respond within the time limit, the trial terminated with a warning message and they needed to press any key to continue. All stimuli were presented in black against the grey background (RGB: 128,128,128). In EEG experiment, the four choices, ‘very real’ (i.e. choosing real with high confidence), ‘slightly real’ (i.e. choosing real with low confidence), ‘slightly pseudo ’ (i.e. choosing pseudo with low confidence) and ‘very pseudo’ (i.e. choosing pseudo with high confidence), were assigned to four keys, ‘s’,’d’,’k’,’l’ on a standard keyboard respectively, and participants were instructed to press the buttons with middle and index fingers of their left and right hands separately. The arrangement of finger pressing was same as that in the EEG experiment but using a MEG-compatible button box. The order of one-to-one match between choice and button was reversed for half of the participants and thus, the presses by left hand and right hand are counter-balanced among participants. Finally, a 1500 ms blank screen appeared following the response. To ensure participants were following the instructions, the catching trials were applied. In catching trials, the stimulus was the label of one of the four choices, such as ‘very real’, instead of a Jurchen character, and participants needed to press the button corresponding to the message. We found catching trials had no priming impact (see supplemental materials for details).

In EEG experiment, participants needed to complete one practice block and four testing blocks. The practice block consisted of 16 trials, including 8 catching trials (the label of each choice was presented twice). Participants needed to pass the practice by responding to 6 or more catching trials correctly out of 8. Each testing block contained 40 testing trials and 3 catching trials. There was a 2-min break between blocks. All stimuli were presented in a random order.

For MEG experiment, the procedure was the same as EEG except that participants needed to complete the practice block twice; one was conducted outside the MEG scanner to familiarize themselves with the task and the other one was conducted under scanning for them to familiarize the scanning environment and the press button. Each block lasted around 6 minutes. To obtain the individual’s brain structure image, after MEG scanning, we required all participants to undergo a structural MRI scan.

### Acquisition and preprocessing of neural data

#### EEG recording and preprocessing

EEG signals were recorded using a Neuroscan Recording System (Neuroscan Inc.,USA) with a 64-channel electrode cap conforming to the standard International 10-20 System of electrode location, sampled at a rate of 1000 Hz. Participants were instructed to minimize masticatory and facial muscular activity, eye movements, and blinking. For measuring the vertical and horizontal electrooculograms (EOG), the EOG signals were recorded from two electrodes placed below one eye and on the external canthus separately. During recoding, all electrodes were online-referenced to right mastoids (M2), and participant’s forehead was connected to the ground through the GND electrode. Impedance of all electrodes was kept below 5 KΩ.

All EEG data were preprocessed with EEGLAB Toolbox (version 14.1.1; Delorme & Makeig, 2004). Raw EEG data were re-referenced to the averaged potentials of bilateral mastoids (M1, M2), down-sampled to 250 Hz, and bandpass-filtered between 0.5 and 40 Hz. These data were epoched into −1300 to 2500 ms relative to the onset of the Jurchen characters, and were corrected with the baseline from −300 to 0 ms. Epochs containing data exceeding 5 standard deviations from the pooled channel mean were removed to optimize ICA (independent component analysis) decomposition, and afterwards the remaining data were decomposed into 62 independent components. Based on the pattern of time domain, frequency and topography of each component, artifactual components (eye blinks, muscle tension, etc.) were detected and then removed (the number of removed components per participant ranged from 3 to 12). Finally, epochs containing voltages exceeding ± 150 μV were excluded. The remaining artifact-free epochs (accepted epochs per participants: Mean = 96.27%, SD = 4.27%) were used for following analyses.

#### MEG recording and preprocessing

MEG signals were recorded using a 306-channel, whole-head magnetometer of Elekta Neuromag (Elekta Oy, Helsinki, Finland), sampled at a rate of 1000 Hz. MEG sensors were arranged in 102 triplets, comprised of one magnetometer and two orthogonal planar gradiometers. Besides, two additional electrodes were placed above the right eye and below the left eye respectively to measure the EOG. In order to facilitate co-registration of the sensor array to participant's individual anatomic MRI of head, prior to MEG scanning, three anatomical fiducial points (nasion, left and right preauricular points) and over 200 additional points evenly spread out over the participant's head (about 10 points digitized at bridge of the nose) were digitized using Probe Position Identification system (Polhemus, VT, USA). During MEG scanning, participants needed to complete 5 blocks, and before each block, the participants’ head position in the MEG helmet was registered by a head position indication system (HPI). Besides, at the beginning of each scanning day, we recorded ~ 2 min empty-room noise data for source reconstruction.

After recordings, the raw data were first preprocessed by temporal Signal Space Separation (tSSS) method. After that, data were preprocessed using the functions from the brainstorm Toolbox (Tadel et al., 2011). Specifically, the offline data were down-sampled to 250 Hz, and high-pass filtered at 0.5 Hz. Then, the data of magnetometers and of gradiometers were separately ICA decomposed, and the top 20 independent components correlated to EOG were extracted. According to the pattern of time domination, topography and frequency for each component, the artifactual components (e.g. heart beats, eye movements, or sensor artifacts) from magnetometer and gradiometers were respectively identified and then removed (the number of removed components per participant ranged from 2 to 5). After removing the artifactual components, data were further inspected with a semiautomatic procedure suggested by brainstorm toolbox to identify the remaining artifacts, and the detected contaminated signals were discarded. Finally, the artifact-free data were epoched into −1300 to 2500 ms relative to the onset of the Jurchen characters and the epoched data were used for the subsequent analyses (accepted epochs per participants: Mean = 93.54%, SD = 2.68%).

#### MRI data acquisition

The structure images of all participants were scanned on a Siemens Prisma 3T MR scanner using a 64-channel head-neck coil. The high resolution anatomical scan was acquired using a 3D Magnetization-Prepared RApid Gradient-Echo (MPRAGE) sequence, 192 continuously sagittal slices, TR = 2530 ms, TE = 2.98 ms, TI = 1100 ms, FA = 7°, bandwidth = 240 Hz/pixel, FOV = 256 mm (FH) × 224 mm (AP), matrix = 256 × 224, slice thickness = 1 mm, voxel size = 1 mm × 1 mm × 1 mm, isotropic, interpolated to 0.5 mm × 0.5 mm × 1 mm, and the time of acquisition = 5 min 58 sec. To reduce the effect of distortion, the distortion of structure images was corrected using Siemens Syngo software.

### Analysis of Behavioral data

#### Choice ratios and reaction times (RTs)

We used two-tailed paired t-test to examine whether the choice ratios and RTs were significantly different in high and low confidence trials. In RT analyses, we excluded the trials in which choice reaction times were below 100 ms or above 3 standard deviations from the mean reaction time per participant (the proportion of excluded trials per participant in MEG: Mean = 2.12%, SD = 2.57%; that in EEG: Mean = 3.71%, SD = 3.68%). Note that, we examined the potential effects of responding hand and Jurchen word-likeliness decision, and found there was no significant ratio and RT difference, either between the left-hand and right-hand presses, or between real choice and pseudo choice trials (see supplemental materials for details).

#### Hierarchical drift-diffusion model parameters

We estimated participants’ drift-diffusion model (DDM) parameters using hierarchical Bayesian estimation with the HDDM toolbox (Wiecki et al., 2013). We fitted the model to the RTs without extreme values (extreme values were removed as RT analyses above) for real and pseudo choices, and estimated the drift rate (v) and decision threshold (a) in high and low confidence trials respectively. In this hierarchical model, individual parameter estimates are constrained by group-level distributions. The model also included inter-trial variability in drift rate (sv) and decision bias (sz and z) as free parameters that were constant across conditions. For the model fitting, we ran the Markov chains each with 10000 samples and the first 1000 of these samples were discarded so as to get more robust results. We inspected the Markov chains and found all chains were convergent.

Here, the upper and lower boundaries were set based on the real choices against pseudo choices instead of on one of four choices (e.g. ‘very real’ choice) against the remaining three. To examine whether the boundary setting biased the DDM results, we conducted another four DDM fitting with each kind of four choices as an upper boundary separately, and the results clearly showed that the fitting parameters of making ‘very real’ choice and that of making ‘very pseudo’ choice were quite similar. The fitting parameters between making ‘slightly real’ choice and making ‘slightly pseudo’ choice were also quite similar (see supplemental materials for details). These findings suggested it was suitable that real and pseudo choices served as upper and lower boundary in one model.

### Analysis of EEG data

For EEG analyses, four participants were excluded. One of them had an extremely low ratio of high confidence choice (less than 2%; the number of trials was smaller than that needs for steady analyses) and another two of them moved head or eyes excessively throughout the recording. The final one was excluded after preprocessing because of the excessive artefact in data that may result from loosening of the online-referenced electrode. Thus, 24 participants (14 females) were included in following EEG analyses. We note the behavioral results of the 24 participants were very similar with those of the 28 participants (i.e. the data with those 4 to-be-excluded participants).

#### Pre-stimulus and post-stimulus neural oscillations at scalp

We examined whether pre-stimulus neural oscillations influenced the decision-confidence in subjective judgment. With an exploratory purpose, we first performed time–frequency analyses from 2 to 45 Hz for all time points (−1.5 to 2.5 s) using a complex Morlet wavelet (central frequency = 1Hz, FWHM =2.7s). We focused on the power of low frequency bands (3 ~ 20Hz) in the period from −1200 to 0 ms to identify pre-stimulus oscillations and that from 0 to 2200 ms to identify post-stimulus oscillations. According to the time–frequency map above and the length of window for steady pre-stimulus time-frequency analyses suggested by previous studies (Iemi et al., 2017; Iemi et al., 2019), the time window from −1000 to −500 ms relative to stimulus onset was selected. To further confirm the power of target frequency bands, the power spectrum density (PSD) was computed for the time window above by Welch’s method (frequency range of 2 to 40 Hz). We noted that the trial number of low confidence choice was overwhelming comparing to that of high confidence choice. Given that the trial number could influence the signal-to-noise ratios and thereby bias the results, we randomly selected largest equal-sized sets of trials for high and low confidence choices per participant and then do contrast analyses based on the selected trials with paired t-test.

#### Binning analyses

We first sorted the trials of each participant according to the pre-stimulus alpha power (i.e. the PSD values at 8 Hz) from low to high and then divided them into four bins evenly. After that, we calculated the proportion of trials with high confidence choice in each bin per participant. Finally, we used Linear Mixed Effects Models (lmerTest function in R) to calculate the relation between high confidence ratios and the pre-stimulus powers, with order of each bin as regressor, bin-level high confidence ratios as dependent variable, and between-participant variance as a random factor.

### Analysis of MEG data

For MEG analyses, four participants were excluded; two of them were due to very low ratios of high confidence choice (i.e. less than 8%; the number of trials was smaller than that needs for steady analyses) and the other two were due to incomplete marker records resulting from a failure in behavioral response synchronizer of MEG recording system. Thus, 26 participants (12 females) were finally included in MEG analyses. We noted the behavioral results of these 26 participants were very similar with those of the 30 participants (i.e. the data with those 4 to-be-excluded participants; see supplemental materials for details).

#### Source reconstruction

The cortex-level source for each time point was localized from MEG data using a constrained weighted minimum-norm estimate (wMNE: depth-weighting factor: 0.5) method built-in brainstorm toolbox (Lin et al., 2006). We produced the overlapping spheres model from 3-D reconstructed MRI with registered MEG sensors for source analysis. When computing the source, the noise covariance for each participant across conditions was computed from the ~ 2 min empty-room data recorded at the same day that participant’s data were collected. Then, the MNE function was called out for whole cortex source computation on MAG and GRAD sensor data, and afterwards the cortex source activation was measured at each time point in the time window from −1300 to 2500 ms.

#### Pre-stimulus and post-stimulus neural oscillations at sensor and cortex level

For the pre-stimulus oscillations, we calculated both the time-frequency map and specific power spectrum density (PSD) map for MEG using the parameters as that for EEG. At sensor level, the time-frequency analyses were conducted with the planar gradiometer representation approach (Ahonen et al., 1993), considering that the transient biological current recorded by EEG channels is very similar to the transient variance of magnetic field recorded by the paired gradiometers, and thereby the gradiometer representation approach makes the EEG and MEG results more comparable. At source level, the PSD was calculated on the whole cortex source of each trial per participant, and then averaged within each participant and condition. These resulting participant-specific source estimates were projected to standard anatomy ICBM152 Nonlinear atlases (Fonov et al., 2011), and went through the 3 mm FWHM (full width at half maxima) spatial smooth. In this study, the pre-stimulus neural oscillation of SMA (MNI: −3.4, −15.1, 57; 32 vertices, 5.04 cm^2^) and MCC/PCC (MNI: 1.2, −14.3, 37.3; 20 vertices, 5.31 cm^2^) were found associated with decision confidence, and thus these two brain areas were set as ROI (region of interesting) for the analyses of post-stimulus neural oscillations. The time-frequency values of ROI were extracted by averaging the time-frequency results from each vertex of the ROI.

#### Phase transfer entropy

Phase transfer entropy (PTE) is a functional connectivity estimation method, which can measure phase-specific directed connectivity from a brain region (e.g. A) to another brain region (e.g. B). Specifically, if A has a causal influence on a target signal of B, the uncertainty of the present of B(t) conditioned on its own past B(t-δ) should be greater than the uncertainty of the present of B(t) conditioned on the signals of both past B(t-δ) and A(t-δ), and TE can be represented by the equation(1) (Lobier et al., 2014).

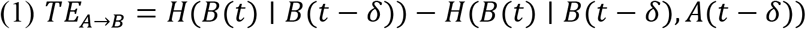

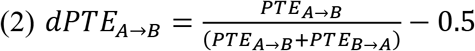

Based on the findings that the pre-stimulus alpha (8Hz) oscillations of both MCC/PCC and SMA enhanced in low confidence trials, we interested in whether the pre-stimulus information flow was directly transferred from MCC/PCC to SMA at 8Hz. To address this, the signals of MCC/PCC and SMA within the time window from −1000 to −500 ms was first time-frequency decomposed with Hilbert transform to get the instantaneous phase time-series in 8Hz. Then, we set MCC/PCC as the brain area A and SMA as the brain area B and used their instantaneous phases in 8Hz to calculate the TE according to the equation (1). Finally, to reduce estimation bias, we applied the equation(2) (Hillebrand et al., 2016) to get the normalized results, dPTE. The values of dPTE ranged from −0.5 to 0.5; thus, positive values indicate that information flowed from PCC/MCC to SMA, while negative values indicate that from SMA to PCC/MCC. The PTE analyses above were achieved also by brainstorm toolbox (Lobier et al., 2014).

#### Binning analyses

The analyses method was the same as that in EEG.

#### Mediation analyses

For mediation analysis, the time window for pre-stimulus PSD of SMA and MCC/PCC was from −1000 to −500 ms, and the windows for post-stimulus PSD of SMA and MCC/PCC were from 1600 to 1900 ms and from 250 to 600 ms respectively. Based on the pre-stimulus alpha powers, we first sorted (from low to high) and divided the trials of each participant into five bins. Then, we calculated the high confidence ratios and the mean of pre-stimulus and post-stimulus PSD values for each bin. We noted that we z-scored the raw PSD within each participants to overcome the raw PSD variance between participants. Finally, we estimated whether the correlation between pre-stimulus PSD and high confidence ratios significantly decreased under statistical control for the post-stimulus PSD following a conventional method (Baron & Kenny, 1986). Specifically, the correlation coefficient between pre-stimulus PSD and high confidence ratios without the statistical control was calculated using a Linear Mixed Effects Model as above. Meanwhile, the correlation coefficient with the control was calculated applying another Linear Mixed Effects Model, using both post-stimulus and pre-stimulus PSD as regressors, bin-level high confidence ratios as dependent variable, and between-participant variance as a random factor.

## Acknowledgements

This work was supported by the Shanghai Leading Talent Plan and the National Natural Science Foundation of China (Grant No. 32071051) to X.Y.G and the National Natural Science Foundation of China (Grant No. 32000770) to L.Z. Z.Y.L. was supported by the National Natural Science Foundation of China (Grant No. 31900754).

## Competing interests

The authors declare that no competing interests exist.

## Supplemental materials

### Behavioral evidence for no reliable linguistic signal from the stimuli materials

As described in the method, the linguistic signals delivered from stimuli were on a level approaching to noise. Here, we rechecked whether participants’ choice was driven by the external linguistic signal of stimulus. If participants’ choices were driven by detecting some external linguistic signal on stimuli, then a subset of stimuli should be consistently rated as real or pseudo with high confidence. The analysis showed no stimulus consistently positioned in the highest or lowest 27% ends of word-likeliness judgment in EEG and MEG experiments. Thus, the external linguistic signal of stimulus has little impact on the participants’ decision.

### The behavioral results of the participants being included in EEG and MEG analyses

Here, we examined whether the behavioral pattern of the participants being included in EEG or MEG analyses was similar to that of all participants. We found, for the participants in EEG analyses, the ratio of low confidence choice (M = 62.78%) was higher than that of high confidence choice (M = 37.23%), *V* = 16, *P* < 0.001. For the participants in MEG analyses, the ratio of low confidence choice (M = 64.04%) was also higher than that of high confidence choice (M = 35.96%), *V* = 13.5, *P* < 0.001. When it comes to RTs, we found, the RT towards the trials of low confidence was longer than that of high confidence for those participants in EEG analyses, *t*(23) = 5.26, *P* < 0.001, Cohen’s d = 1.07, and those in MEG analyses, *t*(25) = 7.35, *P* < 0.001, Cohen’s d = 1.44. The ratios of high confidence choice (EEG: 37.22 %; MEG: 35.96%) were also very similar with that of the previous study (38.10%; Woestmann et al., 2019).

Finally, we fitted DDM to behavioral data for each participant in EEG or MEG analyses, and estimated parameters of drift rate and threshold in high and low confidence trials respectively. The paired t tests on the thresholds showed significant difference between high and low confidence trials for participants in EEG analyses, *t*(23) = 5.48, *P* < 0.001, Cohen’s d = 1.12, and participants in MEG analyses, *t*(25) = 5.76, *P* < 0.001, Cohen’s d = 1.13. However, the difference of drift rates between high and low confidence trials were not significant for either participants in EEG analyses, *t*(23) = 1.07, *P* = 0.295, or participants in MEG analyses, *t*(25) = 1.86, *P* = 0.075. Taken together, the behavioral results of the participants included in EEG or MEG analyses were very similar to those based on the data of all participants.

### The priming effect check for catching trials

Considering there was no obvious external cue related to the judgement task, catching trials here might prime the participants’ choices. To address the question, we examined whether the participant’s choice of the trial next to the catching trial (T+1) was the same as the response instructed by the catching trial (T). The result showed, in trials next to the catching trials (T+1), the proportion that participants gave the same responses as that instructed by the catching trials (T) was 21.11% and 19.6% in EEG and MEG experiments respectively. The ratios were not different than the chance level (25%), *t*(29) = 1.92, *P* = 0.968, in MEG experiment or, *t*(27) = 2.71, *P* = 0.994, in EEG experiment. Thus, there was no priming effect of catching trials.

### DDM estimates for each kind of four choices

Here we examined whether setting real and pseudo choice as the upper and lower boundaries was suitable. To address this, we fitted four DDMs to data with each kind of four choices as upper boundary and the remaining three choices as lower boundary, and extract posterior estimates of DDM parameters for each fitting per participant. Then, we conducted a 2 (confidence: low vs high) ×2 (word-likeness: real vs pseudo) repeated measure ANOVA on the two estimated parameters, drift rate and threshold respectively. The results showed, in EEG experiment, the significant main effect of confidence was measured on threshold, *F*(1,27) = 25.13, *P* < 0.001, partial *η*^2^ = 0.48 and drift rate *F*(1,27) = 26.33, *P* < 0.001, partial *η*^2^ = 0.49, all other effects were not significant, all *F* < 1.27, *P* > 0.271. In MEG experiment, we still only observed main effect of confidence on threshold, *F*(1,29) = 37.59, *P* < 0.001, partial *η*^2^ = 0.56, and drift rate *F*(1,29) = 28.51, *P* < 0.001, partial *η*^2^ = 0.50 (other effects, all *F* < 1.59, *P* > 0.218). The results suggested in both EEG and MEG experiments, the decision processing patterns of real and pseudo choices were very similar while those of high and low confidence choices were very different (Fig. S2). Thus, the results indicated our boundary setting was suitable.

### Examining potential confounding factors in behavioral results

We examined whether the responding hand influenced the participants’ choices and RTs. The results showed, in EEG experiment, there was no significant difference in RTs and choice ratios between left and right hand conditions, both *t* < 1.52, both *P* > 0.141. In MEG experiment, there were no significant responding hand effects either, both *t* < 1.12, *P* > 0.273.

We also examined whether RTs were different between real and pseudo choices, and we did not find significant difference between real and pseudo choice trials either in EEG experiment, *t*(27) = 0.25, *P* = 0.806, or in MEG experiment, t(29) = 1.31, *P* = 0.202.

**Figure S1.**
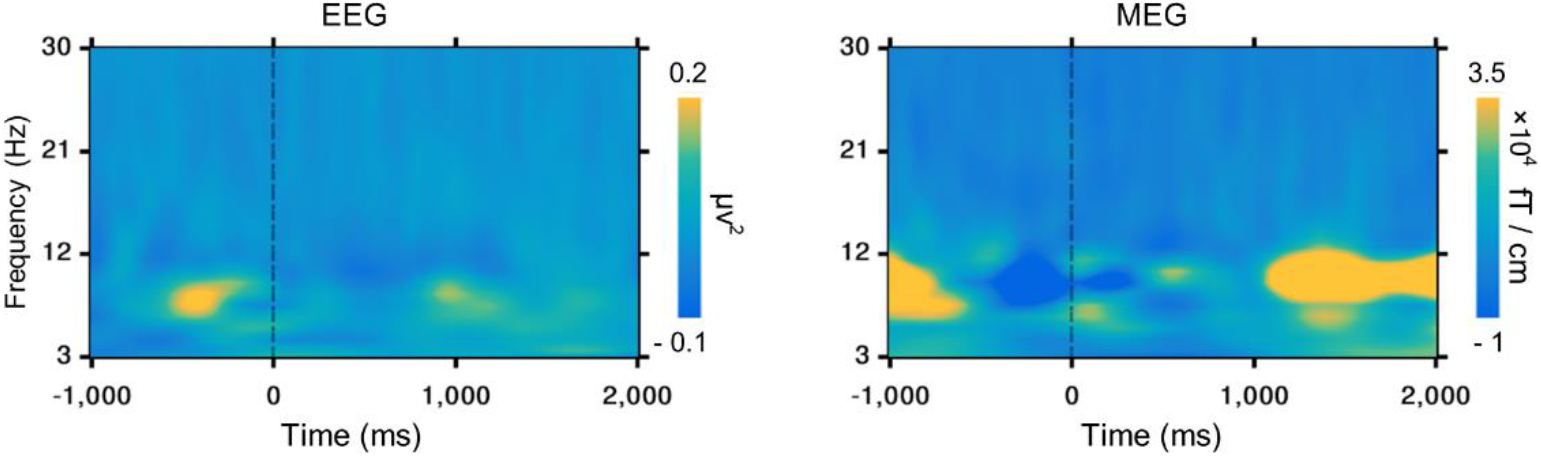
Time-frequency contrast map of low vs. high confidence trials. In both EEG and MEG experiments, stronger pre-stimulus oscillations related to confidence were observed at around 9 Hz. T = 0 indicates the Jurchen character onset.

**Figure S2.**
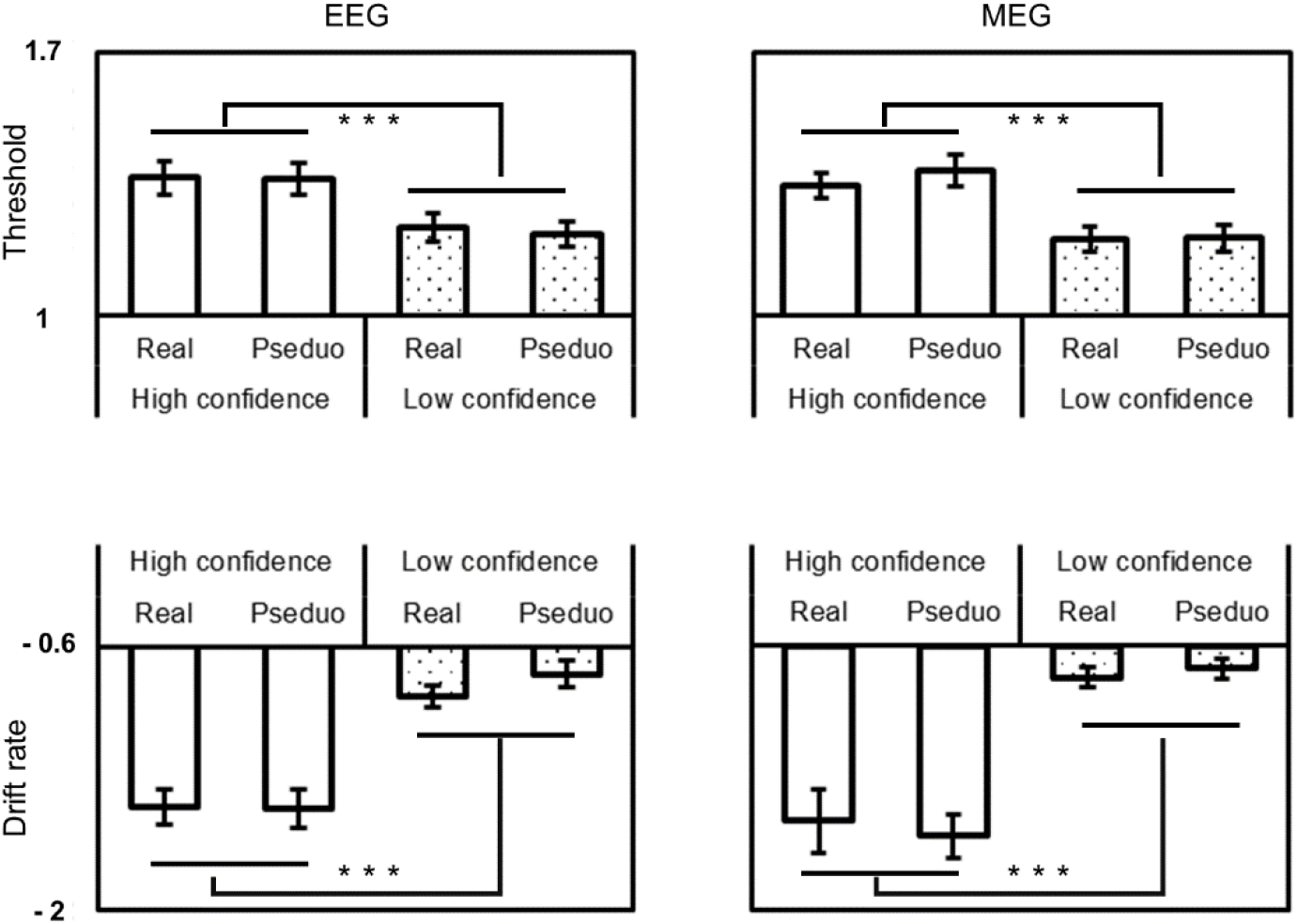
The drift diffusion model fittings for each kind of choices. Here, we fitted four DDMs to data with each kind of four choices as upper boundary and the remaining three choices as lower boundary. In both EEG (left) and MEG (right) experiments, the fitting thresholds (upper) and drift rates (bottom) between real choice trials and pseudo choice trials were quite similar, whereas the parameters between low confidence and high confidence choice trials were significantly different. \*\*\**P* < 0.01. Error bars represent s.e.m.

## Notes

### Competing Interest Statement

The authors have declared no competing interest.

